# Vectofusin-1–based T-cell transduction approach compared with RetroNectin-based transduction for generating murine chimeric antigen receptor T-cells

**DOI:** 10.1101/2021.08.28.458011

**Authors:** Romil Patel, Kartik Devashish, Shubhra Singh, Pranay Nath, Dev Gohel, Rishika Prasad, Manisha Singh, Neeraj Saini

## Abstract

Gene transfer into human and murine T-cells using viral-based approaches has several promising therapeutic applications including the production of chimeric antigen receptor T-cell (CAR-T) therapy. The generation of murine CAR-T is paramount to test and validate immunocompetent mouse models for CAR-T therapy. Several viral transduction enhancers already exist for gene therapy with few limitations. In this study, we tested vectofusin-1, a short cationic peptide, as a soluble transduction enhancer for gammaretroviral transduction for the generation of anti-CD19 murine CAR-T. We found that in comparison to Retronectin, Vectofusin-1 is an equally optimal transduction enhancer for the generation of murine CAR-T cells.

## Introduction

Chimeric antigen receptors (CARs) are synthetic receptors containing a target binding domain usually derived from a single-chain variable fragment of an antibody, a hinge/transmembrane region, and a truncated CD3 zeta cytoplasmic domain with or without a costimulatory domain^1^. CAR T-cell therapy represents the latest advance in the treatment of hematologic malignancies, with unprecedented response rates and survival outcomes seen in patients with relapsed refractory lymphomas^2^. Therefore, there is a need for production of CAR T-cells for both clinical use and laboratory research, particularly mouse validation studies.

Generation of CAR T-cells through viral transduction can be optimized by adding various culture additives such as cationic polymers (polybrene)^3^, dextran^4^, or cationic lipids (lipofectamine^5^). RetroNectin, derived from fibronectin, has recently been used to generate CAR T-cells for clinical applications^6,7^, and RetroNectin has primarily been used to generate mouse T-cells for in vivo testing. However, RetroNectin-based transduction protocols are cumbersome because RetroNectin must be surface-coated prior to use, and therefore, a new soluble additive capable of enhancing infection is needed.

Vectofusin-1, a new cationic amphipathic peptide, is a soluble additive that has been successfully used for transduction of human T-cells; however, this approach has not been tested for generation of murine CAR T-cells. The goal of the current study was to determine whether a novel conjunction protein peptide, such as Vectofuscin-1, can achieve a similar transduction efficiency to that of RetroNectin for generation of CAR T-cells in mice. We have compared the Vectofusin-1–based approach with a more traditional RetroNectin protocol, providing a step-by-step approach to generate murine CAR T-cells using both approaches.

## Materials and Equipment

- RetroNectin (Takara Bio, Catalog No. T100B)
- Vectofusin-1 (Miltenyi Biotec, Catalog No. 130-111-163)
- EasySep Mouse T-Cell Isolation Kit (StemCell, Catalog No. 19851)
- Purified anti-mouse CD3ε antibody (Biolegend, Catalog No. 100302)
- Purified anti-mouse CD28 antibody (Biolegend, Catalog No. 102102)
- EasySep Buffer (StemCell, Catalog No. 20144)
- Non–tissue culture treated 6-well plates
- C57Bl6 mice
- Complete RPMI (cRPMI): RPMI 1640 medium, 10% fetal bovine serum, 100 IU/mL penicillin, 1μM sodium pyruvate, 10mM HEPES, 2.5μM β-mercaptoethanol (added fresh when changing medium), and 2mM L-glutamine
- Phosphate-buffered saline–bovine serum albumin (PBS–BSA): PBS, 0.5% BSA or FACS buffer (PBS + 2% fetal bovine serum)
- Magnet: we used the EasySep magnet, but others are suitable
- Recombinant murine interleukin (IL)-2 (Peprotech, Catalog No. 212-12)
- Frozen viral supernatant from transduced Phoenix-E cells that produce m1928z retrovirus.

## Methods

### Day 1: Mouse T-cell isolation and activation

Note: Mouse spleens can be frozen in fetal bovine serum + 5% DMSO for future use as needed.

#### T-cell isolation

1. Kill mouse and saturate with 70% ethanol. Make a left lateral incision below the rib cage and collect the spleen. Store spleens in cRPMI.
2. Place the spleen onto a 40μM cell strainer placed onto a 50-mL conical tube loaded with 10 mL cRPMI. Using the plunger end of a 1-mL syringe, gently push the spleen through the strainer into the 50-mL tube. Rinse the cell strainer with 10 mL of cRPMI to ensure all spleen cells pass through the strainer into the conical tube.
3. Spin cells at 1500 rpm for 5 minutes, discard supernatant, and wash the pellet once with 5 mL PBS.
4. Resuspend pellet in 5 mL red blood cell lysis buffer.
5. Incubate at room temperature for 2 minutes.
6. Stop red blood cell lysis with 20 mL of cRPMI. Centrifuge and wash once with PBS.
7. Count cells and prepare cell suspension at a concentration of 1 × 10^8^ cells/mL in EasySep buffer.
8. Add normal rat serum from the EasySep Mouse T-cell Isolation Kit (StemCell Technologies) and Isolation Cocktail, both at 50 μL/mL of splenocytes.
9. Mix well and incubate at room temperature for 10 minutes.
10. Vortex EasySep Streptavidin RapidSpheres for 30 seconds and add to the antibody-splenocyte mixture at 75 μL/mL of splenocytes.
11. Mix well and incubate at room temperature for 2.5 minutes.
12. Add EasySep buffer to the splenocyte suspension to a total volume of 2.5 mL and mix by gently pipetting up and down.
13. Place the tube into a magnet and set aside at room temperature for 2.5 minutes.
14. Pick up the magnet, invert, and pour off desired unbound T-cell fraction into a new tube.
15. Spin T-cells at 1500 rpm for 5 minutes, discard supernatant, and resuspend pellet in cRPMI to a final concentration of 1 × 10^7^ cells/mL.

#### T-cell activation

1. Incubate anti-CD3 antibody at 5 μg/mL and anti-CD28 antibody at 2 μg/mL on a 6-well plate (5 mL/well) for 2 hours at 37 °C or overnight at 4 °C (seal the plate). Note: Dynabeads Mouse T-Activator CD3/CD28 (Catalog No. 11456D) can also be used for mouse T-cell activation and expansion, with similar results and expansion.
2. Resuspend T-cells in cRPMI to a final concentration of 1 × 10^7^ cells/mL.
3. Aspirate monoclonal antibody solution from the plates and add 1 mL of T-cells per well.
4. Let cells grow overnight at 37 °C and 5% CO_2_.
5. For transduction with RetroNectin, prepare non-tissue culture treated plates by adding RetroNectin (1 μg/μL; Takara Bio, Otsu, Japan) to enhance transduction efficiency. Add 90 μL of RetroNectin with 6 mL PBS. Dispense 1 mL/well RetroNectin/PBS into non-tissue culture treated plates. Store the plates overnight at 4 °C. Note: The number of wells to be plated with RetroNectin depends on the number of T-cells harvested from the spleen. We prefer to use 1 × 10^7^ cells per well in 6-well plates. RetroNectin-coated plates can also be prepared on the day of transduction, but this requires incubation at room temperature for 2 hours, followed by blocking with PBS-BSA for 30 minutes and washing once with PBS before same-day use.

### Day 2: First transduction

Thaw the recombinant retrovirus supernatant in a 37 °C water bath and remove it from the bath immediately when thawed. Production of gamma-retrovirus production is detailed in the Notes section below.

#### RetroNectin-based transduction

1. Remove RetroNectin/PBS and block with PBS–BSA for 30 minutes at room temperature.
2. Remove blocking buffer and wash plates once with PBS.
3. Add 3 mL of the viral supernatant to RetroNectin-coated wells.
4. Spin at 2000 × g for 1 hour at room temperature.
5. Harvest and centrifuge activated T-cells at 1500 rpm for 5 minutes.
6. Resuspend cells with cRPMI at 1 × 10^7^ cells/mL and 80 IU/mL IL-2.
7. Distribute 1 mL of T-cell suspension to each well.
8. Spin plates at 2000 × g for 1 hour at room temperature.
9. Incubate plates in a tissue culture incubator at 37 °C (5% CO_2_).

#### Vectofusin-1–based transduction (adapted from manufacturer’s instructions)

1. Thaw stock aliquot of Vectofusin-1 (1 mg/mL stock concentration) at room temperature. Vortex thoroughly before use.

a. The required final concentration of Vectofusin-1 for transduction is 10 μg/mL in the total culture volume.
2. Add 10 μL of Vectofusin-1 to 3 mL of viral supernatant and pipette up and down. Note: we have used lower doses of Vectofusin-1, up to 10 μL, without compromising the transduction efficiency.
3. Immediately (within 10 minutes) add the mixture of viral supernatant and Vectofusin-1 to 1 mL of cell suspension (1 × 10^7^ cells/mL) and pipette up and down.
4. Spin plates at 2000 × g for 2 hours at room temperature.
5. Incubate at 37 °C (5% CO_2_).

a. To reach a higher transduction performance, centrifuge cell samples at 400 × g for 2 hours at 32 °C followed by static incubation at 37 °C.

### Day 3: Second transduction

#### RetroNectin transduction

1. Tilt plates and carefully remove most of media from each well, but be careful to not aspirate cells bound to plates at the bottom.
2. Add 3 mL of virus to each well.
3. Spin plates at 2000 × g for 2 hours at room temperature.
4. Incubate in a tissue culture incubator at 37 °C (5% CO_2_).

#### Vectofusin-1 transduction

1. Add 10 μL of Vectofusin-1 to 3 mL of viral supernatant and pipette up and down.
2. Tilt plates and carefully remove most of the media from each well, but be careful to not aspirate cells bound to plates at the bottom.
3. Immediately (within 10 minutes) add the mixture of viral supernatant and Vectofusin-1 to each well and pipette up and down.
4. Spin plates at 2000 × g for 2 hours at room temperature.
5. Incubate at 37 °C (5% CO_2_).

### Day 4-7: Expansion and analysis flow cytometry

1. Collect cells by thoroughly pipetting up and down, spin at 1500 rpm for 5 minutes, and resuspend in 5 mL of cRPMI with 50 IU/mL IL-2 in a T25 flask.
2. Incubate in a tissue culture incubator at 37 °C (5% CO_2_).
3. T-cells can be expanded for up to 7 days without further activation. Fresh culture media can be added or exchanged if the media becomes yellow or the concentration of cells is more than 2 × 10^6^ cells/mL.

### Transduction efficiency analysis

1. Harvest and centrifuge transduced T-cells at 1500 rpm for 5 minutes. Warm cRPMI before thawing frozen T-cells.
2. Wash once with FACS buffer.
3. Stain with Live/Dead dye and stain with the other appropriate markers: CD4, CD8, CD3, CD45.
4. After washing twice, cells can be immediately analyzed using flow cytometry.

### Notes

1. Activation beads can also be used for activation of T-cells; however, they are costly compared with coated antibody methods. We followed the manufacturer protocol, with a 1:1 bead-to-cell ratio. Also, activation beads must be removed from T-cells before analysis or experimental use by pipetting T-cells in a conical tube and thoroughly pipetting up and down to remove the T-cells sticking to beads. If the tube is exposed to a magnet for 2 minutes, the unbound T-cells can be decanted from the tube.
2. There are various protocols for transient production of gamma retrovirus from an ecotropic cell line. However, for large quantities of virus with similar titers, we recommend production of stable virus-producing cell lines, as reported by Li et al^8^. For our experiments, we obtained stable virus-producing Phoenix-Eco cell lines as a generous gift from Dr. Marco L. Davila’s laboratory to generate gamma retrovirus.
3. Human IL-2 can be used for both mouse and human T-cells.

### Statistical analysis

A nonparametric two-tailed *t* test was used to evaluate variation between groups. Differences in proportions were evaluated using a two-sided Fisher exact test. For all statistical analyses, p ≤ 0.05 was considered statistically significant.

## Results

### Transduction efficiency with activation beads compared with plated antibodies

First, we sought to evaluate the activation of murine T-cells and transduction efficiency with beads compared with plated antibodies. We harvested mouse spleens (n = 3) and murine T-cells were isolated as detailed in the protocol above. 5X10^6^ T-cells were collected and were individually activated with either the mouse T-cells activator beads or plated anti-CD3/CD28 antibodies. Subsequently, we performed RetroNectin-based transduction assays with same amount of harvested viral supernatants (3 mL) in each well. Because viral supernatant was obtained from stable virus-producing cell lines and the supernatant was pooled before transduction, we expected the viral titers to be similar between individual experiments and steps.

Our CAR plasmid vector is based on an SFG plasmid backbone with an anti-mouse CD19 CAR sequence followed by a T2A and green fluorescent protein(GFP) sequence, as shown in Figure 1A. We evaluated the transduction efficiency in murine T cells by determining the expression of GFP using flow cytometry on a FITC channel. We found that in RetroNectin-based transduction assays, similar transduction efficiency was generated with beads (mean = 60.40%, n = 3) as with plated anti-CD3/CD28 antibodies (mean = 55.4%, n = 3) using the T-cell activation methods shown in Figure 1B and 1C (p = 0.14, two-tailed unpaired *t* test).

**Figure 1.**
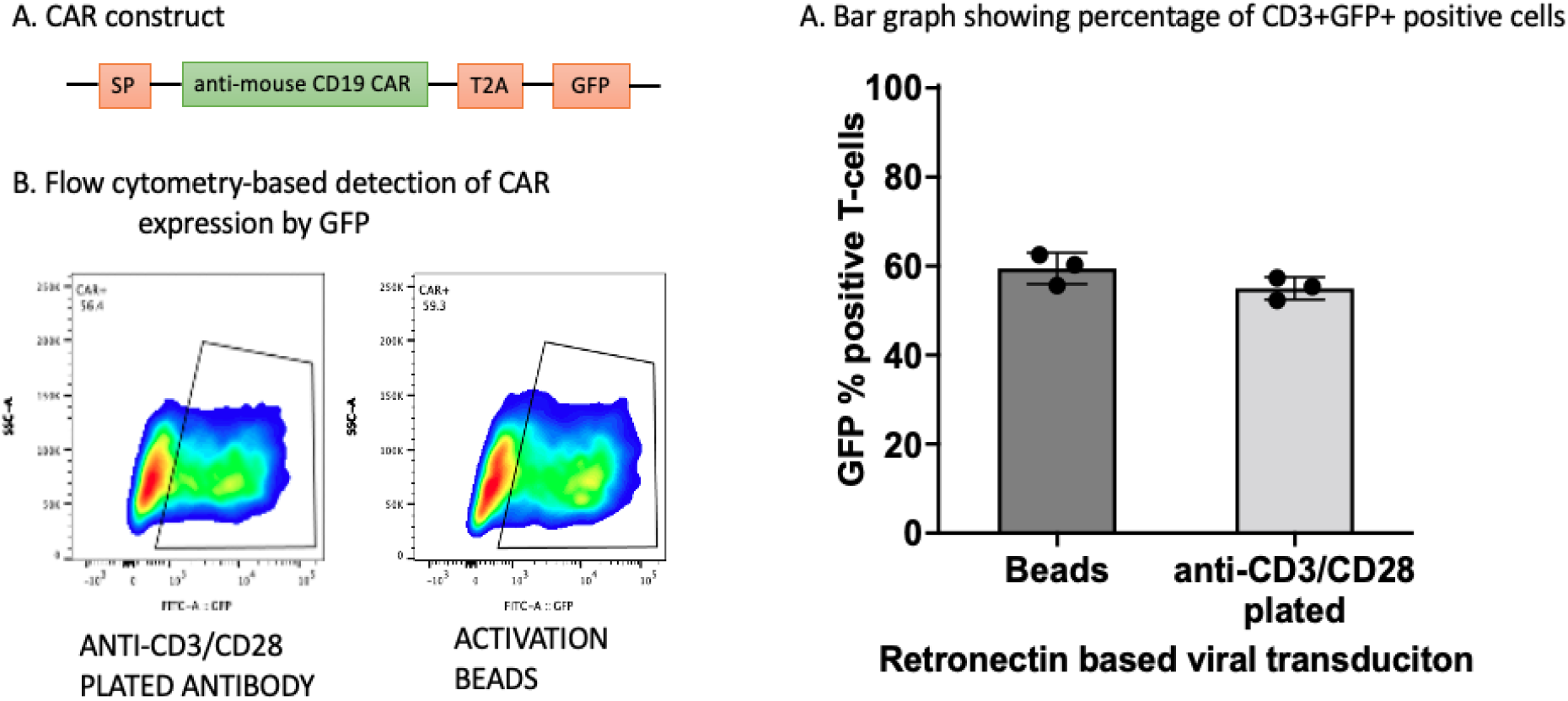
Comparison of anti-CD3/CD28 plated antibodies with a bead-based activation method for generation of murine anti-CD19 chimeric antigen receptor (CAR) T-cells. **(A)** Pictorial representation of CAR vector with a green fluorescent protein (GFP) segment inserted after a flanking T2A sequence. **(B)** Flow cytometry detection of CAR transduction efficiency for bead-based and antibody-based activation. **(C)** Bar graph showing percentages of CD3+GFP+ cells in RetroNectin-based viral transduction assays using beads or plated antibodies for activation.

### Comparison of RetroNectin and Vectofusin-1 for generation of murine CAR T-cells

Next, we compared RetroNectin with Vectofusin-1 as a transduction enhancer for generation of murine CAR T-cells. For direct comparison, we activated T-cells isolated from mouse spleens (n = 3) with plated anti-CD3/CD28 antibodies and did viral transduction on day 1 and day 2. We found that Vectofusin-1 generated lower transduction efficiency (mean = 43.33%, n = 3) than RetroNectin (mean = 55.57%, n = 3; p = 0.001, paired two-tailed *t* test; Figure 2).

**Figure 2.**
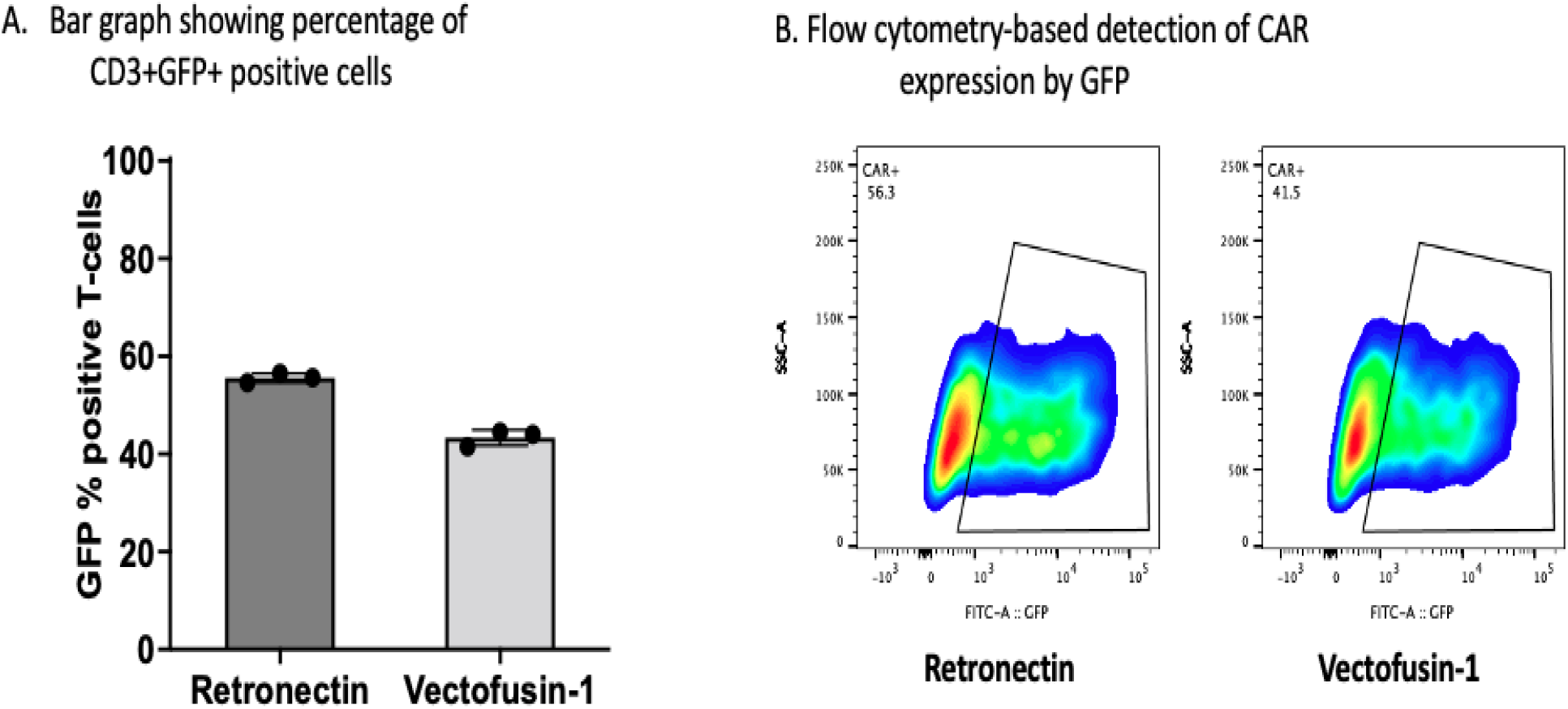
Comparison of Vectofusin-1 with RetroNectin as transduction enhancers for generation of murine chimeric antigen receptor (CAR) T-cells. **(A)** Bar graph showing percentages of green fluorescent protein (GFP)-expressing T-cells. **(B)** Flow cytometry representation of CAR transduction efficiency with RetroNectin and Vectofusin-1.

### CAR-T cells proliferation and expansion is optimal with Vectofusin-1

Next, we also evaluated the absolute number of cells at day 5 after transduction with the help of Countess FL automated Cell Counters and computed the median fold expansion from the baseline. We found that by the end of day 5, fold expansion of total T-cells were significantly higher with Vectofusin-1 (1.9333 + 0.141 fold) compared to Retronectin (1.4667 +0.0706) transduction (p=0.008).

## Discussion

In our study, we compared Vectofusin-1 and RetroNectin as transduction enhancers for the generation of murine anti-CD19 CAR T-cells. We found that the vectofusin-1 generated lesser CAR-T transduction efficiency, as assessed by GFP expression, compared to Retronectin. However, the absolute number of CAR-T cells generated was approximately equal between the two groups due to higher expansion and proliferation of total T-cells under vectofusin-1. The reason for the increased total number of T-cells generated with vectofusin-1 is unclear to us. It is possible that Retronectin labeled plates adhere to T-cells more tightly after spin down post-transduction and probably limit their mobility and proliferation to a small extent. In conclusion, vectofusin-1 leads to optimal murine CAR-T production, like Retronectin, for performing in-vitro and in-vivo mouse validation studies.

## Acknowledgements

We thank Erica Goodoff, Senior Scientific Editor in the Research Medical Library at The University of Texas MD Anderson Cancer Center, for editing this article.

